# Differential inclusion formation of an aggregation-prone protein reveals differences in the proteostasis capacity of neuronal cell lines

**DOI:** 10.1101/2025.08.26.672484

**Authors:** Shannon McMahon, Dezerae Cox, Flora Cheng, Albert Lee, Justin J. Yerbury, Heath Ecroyd

**Affiliations:** Molecular Horizons and School of Science, University of Wollongong, Wollongong, Australia; Yusuf Hamied Department of Chemistry, University of Cambridge, Cambridge, UK; UK Dementia Research Institute, University of Cambridge, Cambridge, UK; Macquarie University Motor Neuron Disease Research Centre, Macquarie University, Sydney, Australia

**Keywords:** Protein aggregation, Inclusions, Proteostasis capacity, Neurodegenerative disorders, Aggregation index

## Abstract

Maintaining proteome integrity is essential for cellular function and survival. Disruptions in proteostasis lead to the aggregation of proteins into inclusions, a process that underlies many neurodegenerative diseases. To quantitatively assess the proteostasis capacity of neuronal cells, we employed an aggregation-prone double mutant form of firefly luciferase (denoted Fluc^DM^) as a reporter protein. We compared two commonly used neuronal cell lines, mouse neuroblastoma cells (Neuro-2a) and a motor neuron-like hybrid line (NSC-34), to evaluate their ability to prevent the aggregation of proteins into intracellular inclusions. We observed a significantly greater propensity of Fluc^DM^ to form inclusions in NSC-34 cells compared to Neuro-2a cells. This suggests a reduced capacity of NSC-34 cells for managing aggregation-prone proteins. Proteomic profiling of Fluc^DM^ inclusions purified from both cell types revealed cell-type-specific engagement of the proteostasis machinery with aggregation-prone proteins. Comparing the proteomic profiles of key arms of the proteostasis network between these two cell lines revealed that the endoplasmic reticulum (ER) unfolded protein response is differentially expressed. This study establishes a quantitative platform for assessing cellular proteostasis capacity and underscores the importance of cell-type context in proteome maintenance. These insights have implications for understanding the selective vulnerability of neurons in protein misfolding disorders.

## Introduction

Under normal cellular conditions, native proteins are in equilibrium with partially-folded intermediate states and completely unfolded (denatured) conformations, as part of the protein on-folding pathway [1]. However, certain circumstances, such as gene mutations, epigenetic factors, aging and cell stress, can result in partially-folded intermediates persisting at higher than normal concentrations, leading to increased exposure of hydrophobic side chains [2]. Exposed hydrophobic regions are attracted to similar hydrophobic surfaces on neighbouring molecules and thus, these partially-folded intermediates are prone to self-association and aggregation [3, 4]. This process results in proteins exiting the protein on-folding pathway and entering off-folding pathways [5, 6], leading to the formation of amorphous (disordered) aggregates or highly ordered amyloid fibrils [7–9]. An evolutionarily conserved action of cells in response to stress is the localisation of misfolded or aggregated proteins into subcellular protein deposits, termed protein inclusions [10–12]. The formation of these proteinaceous inclusions is a pathological hallmark of many debilitating conditions, some of the most notable being neurodegenerative diseases such as Alzheimer’s diseases, Parkinson’s disease and amyotrophic lateral sclerosis (ALS).

Different cell types have varying susceptibilities to inclusion formation [13–16]. For example, inclusion formation in neurons and glial cells is associated with neurodegenerative disease, despite many aggregation-prone proteins being expressed (sometimes at higher concentrations) throughout the body [17, 18]. However, the molecular basis of this susceptibility to inclusion formation remains unknown. The innate ability to maintain aggregation-prone proteins in a soluble state is known to vary significantly between cell types [15, 19]. Neurons are a primary example of this, having a limited capacity to dilute toxic protein aggregates via cell division, stemming from their increased longevity compared to other cell types [19–21]. Furthermore, the transcriptome, proteome and metabolome have also been shown to be characteristic of specific cell types [22]. Thus, intrinsic differences in the pathways that regulate the intracellular environment may also explain the susceptibility of different cell types to inclusion formation.

Protein homeostasis, or proteostasis, refers to the maintenance of the conformational and functional integrity of the proteome. The term proteostasis therefore encompasses all of the pathways that regulate the synthesis, concentration, folding, trafficking and degradation of proteins. The proteostasis capacity of a cell can be defined as the ability to prevent protein accumulation and aggregation into intracellular inclusions. Thus, proteostasis capacity is highly dependent on the protein quality control network. The protein quality control network comprises the systems that act to prevent protein aggregation, for example, through proper guidance of protein folding or assisting in protein degradation [23]. These systems encompass all of the responses in the cell that are activated by proteotoxic stress and include the chaperome, endoplasmic reticulum (ER) unfolded protein response, the heat shock factor 1 (HSF1)-mediated heat-shock response and proteolytic processing by the degradation machinery (i.e. autophagy or the ubiquitin-proteasome system). It has been hypothesised that the relative susceptibility of some cells to inclusion formation is due to intrinsic differences in the cellular protein quality control systems that maintain proteostasis [24–29].

In order to better understand the molecular mechanisms that underpin diseases associated with protein aggregation, and to advance the development of therapeutic strategies, methods to quantitatively measure the proteostasis capacity of a cell are essential. This is especially true given that the proteostasis capacity of a cell is known to influence the rate at which misfolded proteins accumulate [30–33]. Thus, a decline in proteostasis capacity is linked to an impaired ability of a cell to prevent protein aggregation, which can lead to the onset and progression of toxicity associated with disease [34, 35]. The ability to measure the proteostasis capacity of different cell types may help us to deduce why some cells are inherently more prone than others to aggregation and associated cell death.

Here, we exploit a double mutant (DM; R188Q, R261Q) aggregation-prone firefly luciferase (Fluc^DM^) protein [36] to quantitatively compare the ability of two neuronal cells lines (mouse neuroblastoma cells [Neuro-2a] cells and mouse neuroblastoma × motor neuron hybrid [NSC-34] cells) to prevent inclusion formation. Previous work conducted to delineate differences in the proteomes of Neuro-2a and NSC-34 cells revealed that these cell types are highly similar with regard to their global proteomes [37]. However, our work demonstrates that Fluc^DM^ forms inclusions more readily in NSC-34 cells compared to Neuro-2a cells. The inclusions formed by Fluc^DM^ in both Neuro-2a and NSC-34 cells were purified in order to identify differences in the proteostasis machinery that engages with aggregation-prone proteins in these cells. By exploiting a model aggregation-prone protein to quantitatively measure the proteostasis capacity of cells, this work provides insights into the vulnerability of some cells to proteome dysfunction, leading to the formation of inclusions.

## Materials and methods

### Plasmids

The pN3-enhanced green fluorescent protein (EGFP) plasmid was donated by Associate Professor Darren Saunders (University of New South Wales, Australia). Plasmids encoding wild-type (WT) and double mutant (DM; R188Q, R261Q) Fluc with an N-terminal EGFP tag (Fluc^WT^-EGFP and Fluc^DM^-EGFP) [36] were gifted by Professor Ulrich Hartl (Max Planck Institute of Biochemistry, Munich, Germany) and were cloned into pcDNA4/TO/myc/hisA for mammalian expression by GenScript (Piscataway, NJ, USA).

### Cell culture, transfection and treatment of Neuro-2a and NSC-34 cells

Neuro-2a and NSC-34 cells (American Type Culture Collection, Manassas, VA, USA) were cultured in DMEM/F-12 supplemented with 2.5 mM L-glutamine (Gibco, Carlsbad, CA, USA) and 10% (v/v) foetal calf serum (FCS) (Gibco) at 37°C under 5% CO_2_/95% air in a Heracell 150i CO_2_ incubator (Thermo Fisher Scientific, Glen Burnie, MD, USA) and were routinely tested for mycoplasma contamination (∼ every 6 months). For transient transfections, 1.3 × 10^5^ cells/mL were seeded in CELLSTAR^®^ 6-well plates (Greiner Bio-One, Frickenhausen, Germany) or 8-well chamber µ-Slides (Ibidi, Martinsried, Germany). In a 6-well plate, cells were transfected 24 h post-plating with 2 μg/well of either EGFP, Fluc^WT^-EGFP or Fluc^DM^-EGFP plasmid DNA using Lipofectamine^®^ LTX/PLUS^™^ (Life Technologies, Carlsbad, CA, USA; 6 μL/well of Lipofectamine^®^ LTX and 2 μL/well PLUS^™^ reagent) or Lipofectamine^™^ 3000 (Life Technologies; 3 μL/well of Lipofectamine^™^ 3000 and 4 μL/well P3000 reagent) reagents, according to the manufacturer’s instructions. DNA/reagent complexes were added in a dropwise manner to cells. Untransfected controls received equivalent volumes of Lipofectamine^®^ LTX/PLUS^™^ or Lipofectamine^™^ 3000 reagents but did not receive any plasmid DNA.

### Confocal microscopy

To assess the inclusions formed following expression of Fluc^WT^-EGFP, Fluc^DM^-EGFP or EGFP (as a control), Neuro-2a and NSC-34 cells were plated in 8-well chamber µ-Slides as above, transfected using one tenth of the amount of plasmid DNA and transfection reagent so as to maintain the same ratio of these reagents in the volume of culture medium as transfections performed in 6-well plates. Cells were incubated at 37°C for 48 h prior to being analysed by confocal microscopy. Live cells were analysed directly in 8-well chamber µ-Slides using a Leica TCS SP5 confocal microscope and the 63× oil-immersion objective lens (Leica Microsystems, Wetzlar, Germany) controlled by the Leica Application Suite (LAS)-X (Leica Microsystems) version 3 software. EGFP fluorescence was detected by excitation at 488 nm.

### Flow cytometric assays

In some experiments, 48 h post-transfection, cells were prepared for flow cytometric analysis. To do so, cells were harvested with 0.05% (v/v) trypsin/EDTA (Gibco), then diluted with DMEM/F-12 containing 1% (v/v) FCS and centrifuged at 300 × *g* for 5 min at RT. Cells were then washed twice in phosphate buffered saline (PBS; 135 mM NaCl, 2.7 mM KCl, 1.75 mM KH_2_PO_4_, 10 mM Na_2_HPO_4_, pH 7.4) and resuspended in 500 μL PBS. Cells were kept on ice throughout this process to minimise cell death.

The cell suspension was taken and analysed by flow cytometry. Flow cytometry was performed using a BD LSRFortessa X-20 analytical flow cytometer (BD Biosciences, San Jose, CA, USA) and FCS files were analysed using FlowJo version 10 (Tree Star Ashland, OR, USA). The percentage of cells containing inclusions was identified by the previously described technique known as PulSA [38], using the excitation wavelength and emission collection window for EGFP (488 nm, 525/50 nm, respectively). In all experiments a minimum of 100,000 events were acquired, unless otherwise specified. Briefly, in addition to fluorescence area, the width and height parameters of the EGFP fluorescence signal of each event were recorded and used to determine the number of cells with inclusions. The PulSA technique facilitates the identification of cells with inclusions as a result of a shift in their EGFP fluorescence profile, such that they have a narrower EGFP fluorescence pulse width and increased EGFP fluorescence pulse height compared to cells expressing EGFP alone. To account for differences in protein expression between the Neuro-2a and NSC-34 cell lines, PulSA analysis was only conducted on an equivalent subset of EGFP-positive events expressed by both cell types.

In order to compare the relative propensity of Fluc to form aggregates in Neuro-2a and NSC-34 cells (and hence the capacity of cells to prevent aggregation), two different strategies were employed involving the use of PulSA. First, live, EGFP-positive cells were identified and selected (using the untransfected cells as a negative control), and the geometric mean of the EGFP fluorescence in EGFP-positive cells was determined (indicating the level of Fluc expression in the cell). The percentage of cells identified to contain inclusions, as assessed by PulSA, along with the EGFP geometric mean (of cells expressing Fluc^WT^-EGFP or Fluc^DM^-EGFP) were used to generate a PulSA aggregation index:

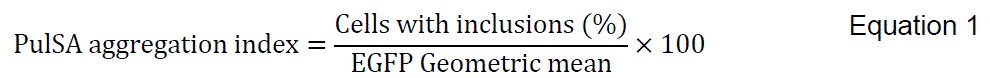

A second approach using PulSA was employed to determine the proportion of cells with inclusions as the levels of Fluc increased in the cells. To do so, EGFP-positive cells were identified and binned into groups based on their levels of EGFP fluorescence: On a log_10_-based scale, 16 gates (bins) of equal size were used to subdivide the EGFP-positive cells. PulSA was then performed on cells within each of these bins to determine the proportion of cells with inclusions in each bin (only bins with >100 cells were analysed). Data are presented as a plot of the bin number versus the proportion of cells with inclusions in that bin.

Histograms were generated and statistical analyses were performed using GraphPad Prism version 8 (GraphPad Software, San Diego, CA, USA). Unless otherwise stated, results are reported as the mean ± standard error of the mean (S.E.M) and the number of independent (biological) replicates (n) of each experiment is specified. Data were analysed by two-way analysis of variance (ANOVA) and Bonferroni’s post-hoc test or, where appropriate, assessed assuming unequal variance using the unpaired two-tailed Student’s t-test. In all analyses, *P* < 0.05 was considered statistically significant.

### Purification of Fluc^DM^ inclusions from SDS-PAGE

In order to purify Fluc^DM^-EGFP inclusions from Neuro-2a and NSC-24 cells to identify proteins contained within these inclusions, transfected cells were trypsinised, harvested, washed twice in PBS (300 × *g* for 5 min at RT) and total cellular protein was extracted by lysis with Nonidet^™^ P-40 (NP-40; Thermo Fisher Scientific) lysis buffer (50 mM Tris-HCl, 150 mM NaCl, 1 mM EDTA, 1% (v/v) NP-40 supplemented with 0.5% (v/v) Halt^™^ Protease and Phosphatase Inhibitor Cocktail (Thermo Fisher Scientific), pH 8.0). Cell lysates were sonicated using the Sonifer^®^ 250 Digital cell disruptor and a double step micro-tip (Branson Ultrasonics, Brookfield, CT, USA) at 50% amplitude for 5 sec. The total protein concentration for each sample was then determined using a BCA assay (Thermo Fisher Scientific) according to the manufacturer’s instruction. The concentration in each sample was adjusted with NP-40 lysis buffer to generate cell lysates of 1 mg/mL total protein (total volume was 200 µL) to ensure equal loading onto SDS polyacrylamide gel electrophoresis (SDS-PAGE) gels. The cell lysate was centrifuged at 20,000 × *g* for 30 min at 4°C and the pellet was washed in ice-cold TNE buffer (50 mM <u>T</u>ris-HCl, 150 mM <u>N</u>aCl, 1 mM <u>E</u>DTA, pH 8.0) and centrifuged again at 20,000 × *g* for 30 min at 4°C. The supernatant was carefully removed and discarded and the pellet resuspended in 50 µL NP-40 lysis buffer. The insoluble pellet was sonicated at 50% amplitude for 5 sec (NP-40 insoluble fraction). Native loading buffer without reducing agents (final concentrations: 200 mM Tris-HCl, 40% (v/v) glycerol, 0.01% (w/v) bromophenol blue, pH 8.6), was added to insoluble cell lysates and the samples were heated at 45°C for 5 min prior to loading onto SDS-PAGE gels. Equal amounts of protein were loaded onto polyacrylamide resolving gels (12% (w/v) acrylamide/bis, 375 mM Tris (pH 8.8), 0.1% (w/v) SDS, 0.25% (v/v) tetramethylethylenediamine, 0.02% (w/v) ammonium persulfate) with polyacrylamide stacking gels (4% (w/v) acrylamide/bis, 330 mM Tris (pH 6.8), 0.1% (w/v) SDS, 0.4% (v/v) tetramethylethylenediamine, 0.04% (w/v) ammonium persulfate) using Precision Plus Protein^™^ dual colour standards (Bio-Rad, Hercules, CA, USA). The gels were run in a Mini-Protean^®^ Tetra Cell system (Bio-Rad) filled with SDS-PAGE running buffer (25 mM Tris base, 192 mM glycine, 0.5% (w/v) SDS, pH 8.3). Samples were electrophoresed at 150 V and were allowed to run until the bromophenol blue dye front had migrated off the end of the gel (∼1 h).

Gels were placed into MilliQ water and immediately imaged using the ChemiDoc^™^ Imaging System (Bio-Rad), and the Pro-Q Emerald 488 exposure setting (exposure time ∼ 5 sec) to visualise EGFP fluorescence from Fluc^DM^-EGFP. Proteins were then fixed in Coomassie Blue staining solution (40% (v/v) methanol, 10% (v/v) acetic acid, 0.02% (w/v) Coomassie Brilliant Blue R-250) and de-stained using de-staining solution (40% (v/v) methanol, 10% (v/v) acetic acid). Regions of the stacking gel identified to contain aggregated Fluc^DM^-EGFP (based on the in-gel fluorescence) were cut out using a clean razor blade and fixed in 10% (v/v) acetic acid for proteomic mass spectrometry. Untransfected Neuro-2a and NSC-34 cells were used as controls and treated as above in order to identify endogenous proteins or protein complexes that are found in the same region of the stacking gel. The subsequent in-gel trypsin digest and proteomic mass spectrometry of the Fluc^DM^ inclusions were performed as described below.

### In-gel trypsin digestion

Excised protein gel bands were further de-stained in 50% (v/v) acetonitrile (ACN) and 50 mM ammonium bicarbonate (pH 8.0) and dehydrated in 100% ACN. The gel pieces were dried by vacuum centrifugation, reduced with 10 mM dithiothreitol at 55°C for 30 min and alkylated with 20 mM iodoacetamide at RT for 30 min. The gel pieces were then rehydrated with 12.5 ng/µL trypsin (Promega, Madison, WI, USA) and resuspended in 50 mM ammonium bicarbonate (pH 8.0), where the proteins were then digested overnight at 37°C. The digestion was inactivated by the addition of 2 μL of formic acid. Tryptic peptides were extracted twice with 50% (v/v) ACN and 2% (v/v) formic acid and dried under vacuum centrifugation. The peptides were resuspended in 0.1% (v/v) formic acid and desalted on a pre-equilibrated C_18_ Omix Tip (Agilent, Santa Clara, CA, USA) and eluted in 50 % (v/v) ACN, 0.1% (v/v) formic acid, and dried under vacuum centrifugation.

### Reverse phase C_18_ liquid chromatography mass spectrometry (RP-LC-MS/MS)

Lyophilised peptides were resuspended in 0.1% (v/v) formic acid and sonicated for 20 min in a sonication bath (Branson Ultrasonics). The resuspended peptides were then centrifuged at 14,000 × *g* for 15 min to remove any insoluble debris, and the clarified peptides were analysed by LC-MS/MS. The peptides were separated on an UltiMate^™^ 3000 RSLCnano system (Thermo Fisher Scientific) fitted with an Acclaim PepMap RSLC column (Thermo Fisher Scientific), making use of a 60 min gradient (2-95% (v/v) ACN, 0.1% (v/v) formic acid) running at a flow rate of 300 nL/min. Peptides eluted from the nano-LC column were subsequently ionised into the Q Exactive^™^ Plus Hybrid Quadrupole-Orbitrap^™^ Mass Spectrometer (Thermo Fisher Scientific). The electrospray source was fitted with a 10 μm emitter tip (New Objective, Woburn, MA, USA) and maintained at 1.6 kV electrospray voltage. The temperature of the capillary was set to 250°C. Precursor ions were selected for MS/MS fragmentation using a data-dependent “Top 10” method operating in Fourier transform (FT) acquisition mode with Higher C-trap Dissociation (HCD) fragmentation. FT-MS analysis on the Q Exactive^™^ Plus was carried out at 70,000 resolution and an automatic gain control (AGC) target of 1×10^6^ ions in full MS. MS/MS scans were carried out at 17,500 resolution with an AGC target of 2×10^4^ ions. Maximum injection times were set to 30 and 50 milliseconds, respectively. The ion selection threshold for triggering MS/MS fragmentation was set to 25,000 counts and an isolation width of 2.0 Da was used to perform HCD fragmentation with normalised collision energy of 27.

Raw spectra files were processed using the Proteome Discoverer software 2.4 (Thermo Fisher Scientific) incorporating the Sequest search algorithm. Peptide identifications were determined using a 20 ppm precursor ion tolerance and a 0.1 Da MS/MS fragment ion tolerance for FT-MS and HCD fragmentation. Carbamidomethylation modification of cysteines was considered a static modification while oxidation of methionine, deamidation of asparagine and glutamine, and acetyl modification on N-terminal residues were set as variable modifications allowing for a maximum of two missed cleavages. The data were processed through Percolator for estimation of false discovery rates. Protein identifications were validated employing a q-value of 0.01. The relative abundance of proteins within each sample was calculated by the Proteome Discoverer 2.4 software using the cumulative intensities of unique peptides. Only proteins identified with an abundance ≥ 2-fold higher than in the untransfected control, and which appeared in all three biological replicates of each sample were considered for further analyses. The mass spectrometry proteomics data have been deposited to the ProteomeXchange via the PRIDE partner repository (PubMed ID: 34723319) with the identifier PXD031826.

### Functional pathway enrichment analysis of proteins identified within Fluc^DM^ inclusions

Venn diagrams constructed to show the overlap of proteins identified in Fluc^DM^ inclusions in Neuro-2a and NSC-34 cells were produced using the Ven de Peer Lab Bioinformatics and Evolutionary Genomics online tool (http://bioinformatics.psb.ugent.be/webtools/Venn). The functional enrichment analysis of the Kyoto Encyclopedia of Genes and Genomes (KEGG) biological pathways represented by proteins identified to co-interact with Fluc^DM^ inclusions in Neuro-2a and NSC-34 cells was conducted using g:Profiler [39] with the following parameters: organism: *Mus Musculus*, statistical domain scope: only annotated genes, multiple testing significance threshold: g:Profiler tailor made g:SCS, user threshold: *P* ≤ 0.05 and data sources: biological pathways – KEGG. To identify the classes of proteins enriched in these samples, Protein Annotation through Evolutionary Relationship (PANTHER) analysis was conducted using the following settings: organism: *Mus Musculus*, analysis: functional classification viewed as pie chart, ontology: protein class. Pie charts were constructed whereby the number of genes was expressed as a percentage of the total number of protein class hits. The Search Tool for the Retrieval of Interacting Genes/Proteins (STRING) analysis [40] of overlapping proteins identified with Fluc^DM^ inclusions in either Neuro-2a or NSC-34 (or both) cells were performed using Cytoscape [41] and data are presented to show how the proteins identified were clustered based on protein class using the Reactome database [42].

### Immunoblotting and detection

Neuro-2a and NSC-34 cells were transfected, harvested and fractionated as above. Samples containing the insoluble fraction were added to SDS-PAGE loading buffer (final concentrations: 500 mM Tris-HCl, 2% (w/v) SDS, 25% (v/v) glycerol, 0.01% (w/v) bromophenol blue, 15% (v/v) β-mercaptoethanol, pH 6.8), heated at 95°C for 5 min and separated by SDS-PAGE as described previously prior to transfer onto a ImmunoBlot^™^ polyvinylidene difluoride (PVDF) membrane (Bio-Rad) using a standard technique [43]. Briefly, proteins were transferred onto a PVDF membrane at 100 V for 1 h in ice-cold transfer buffer (25 mM Tris base, 192 mM glycine, 20% (v/v) methanol, pH 8.3). The membrane was blocked at 4°C overnight with 5% (w/v) skim milk powder in Tris-buffered saline (TBS; 50 mM Tris base, 150 mM NaCl, pH 7.6). Membranes were incubated with primary antibodies of interest [DnaJB6 (1:1000; sc-365574, Santa Cruz Biotechnology, Dallas, TX, USA), Hsp90 (1:1000; ab13492, Abcam) or Hsp105 (1:1000; sc-74550, Santa Cruz Biotechnology)] in 5% (w/v) skim milk powder in TBS containing 0.05% (v/v) Tween 20 (TBS-T) for 2 h at RT. The membrane was washed four times (each for 10 min) in TBS-T before being incubated with either a goat anti-rabbit IgG horse radish peroxidase (HRP)-conjugated antibody (Thermo Fisher Scientific; 31466) or a rabbit anti-mouse IgG HRP-conjugated antibody (Sigma-Aldrich; A9044), diluted 1:5000 into 5% (w/v) skim milk powder in TBS-T. The membrane was rocked at RT for 1 h before being washed four times (each for 10 min) in TBS-T. Proteins of interest were detected with SuperSignal^™^ West Dura Extended Duration Substrate or SuperSignal^™^ West Femto Maximum Sensitivity Substrate (Thermo Fisher Scientific) using an Amersham Imager 600RGB (GE Healthcare Life Sciences, Little Chalfont, UK) or ChemiDoc^™^ Imaging System, with exposure times ranging from 1–5 min.

### Gene ontology mapping of proteins in Neuro-2a and NSC-34 cells

Targeted gene ontology (GO) term analysis was performed using custom python scripts available via Zenodo (DOI: 10.5281/zenodo.16892326), using a keyword-based filtering approach. First, log_2_-transformed abundance values for individual proteins quantified in both the Neuro-2a and NSC-34 cell lines were extracted from the previously published dataset [37], retaining only entries with a unique, reviewed UniProt accession number. Proteins were then grouped based on their annotation with GO biological process terms containing specific keywords of interest. The search terms included: (1) “chaperones”, (2) “ubiquitin” “proteasome” system, (3) “autophagy”, (4) “unfolded protein” response, or the specific term (5) “GO:0030968 (ER unfolded protein response)”. The relative expression of proteins in each group were then analysed directly via linear regression, or via the creation of an abundance ratio (Neuro-2a/NSC-34). The distribution of the abundance ratios for each group was compared to that of the global proteome using a non-parametric Wilcoxon rank-sum test to identify groups whose abundance profiles differed significantly from the whole proteome.

## Results

### Expression of aggregation-prone Fluc^DM^ uncovers enhanced vulnerability of NSC-34 cells to inclusion formation compared to Neuro-2a cells

The aggregation propensity of Fluc^DM^ was exploited in order to measure and compare the relative capacities of Neuro-2a and NSC-34 cells to prevent inclusion formation by aggregation-prone proteins. First, the presence and localisation of inclusions formed by Fluc in Neuro-2a (*top*) and NSC-34 (*bottom*) cells was analysed 48 h post-transfection by confocal microscopy (Fig. 1). Neuro-2a and NSC-34 cells expressing EGFP (*left*) exhibited soluble and diffuse green fluorescence throughout the cytoplasm and the nucleus. Punctate inclusions formed by Fluc^WT^-EGFP (*centre*) were occasionally observed in transfected Neuro-2a and NSC-34 cells. Both Neuro-2a and NSC-34 cells expressing Fluc^DM^-EGFP (*right*) had a greater proportion of transfected cells containing inclusions compared to cells expressing EGFP or Fluc^WT^-EGFP. Overall, it was concluded that Fluc^DM^-EGFP forms inclusions in both Neuro-2a and NSC-34 cells under these experimental conditions.

**Fig. 1.**
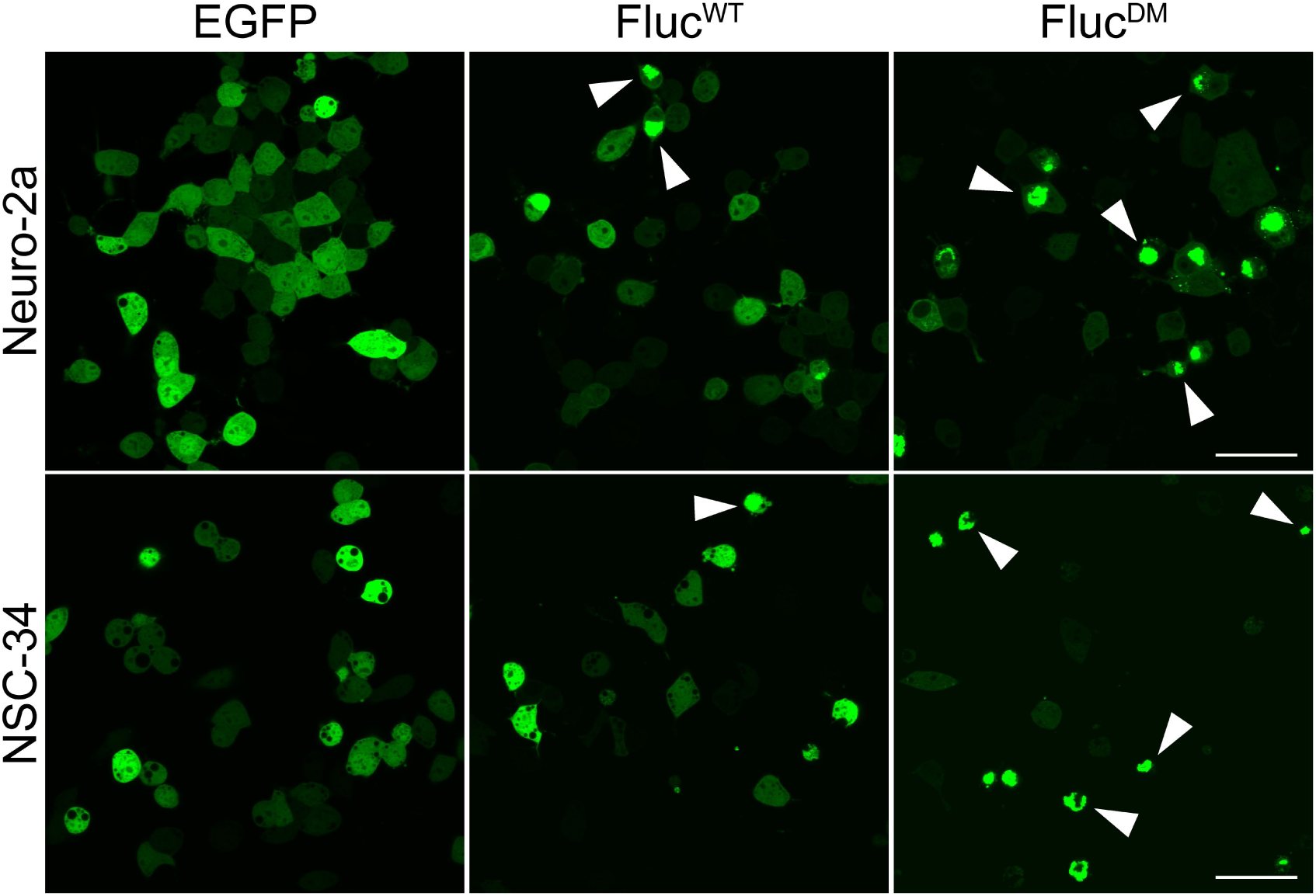
Formation of inclusions by Fluc in Neuro-2a and NSC-34 cells, as assessed by confocal microscopy. Neuro-2a and NSC-34 cells were transiently transfected to express EGFP, Fluc^WT^-EGFP or Fluc^DM^-EGFP and live cells were analysed 48 h post-transfection by confocal microscopy to examine the formation of inclusions. Images are representative of three independent experiments and show confocal micrographs of Neuro-2a (*top*) and NSC-34 (*bottom*) cells transfected with EGFP (*left*), Fluc^WT^-EGFP (*centre*) or Fluc^DM^-EGFP (*right*). Arrowheads indicate cells with inclusions. Scale bar represents 50 µm

Next, we sought to determine the relative capacity of Neuro-2a and NSC-34 cells to maintain Fluc^DM^-EGFP in a non-aggregated state. Cells were transiently transfected with EGFP, Fluc^WT^-EGFP or Fluc^DM^-EGFP for 48 h and then analysed via the flow cytometric method pulse shape analysis (PulSA). Polygonal gating of the forward scatter (FSC) and side scatter (SSC) signals was used to identify viable, live cells and to remove dead cells, cellular debris and cell doublets from subsequent analyses (Fig. 2A). Transfected cells were selected based upon EGFP fluorescence, using untransfected cells as a control; cells that did not fluoresce were excluded from further analyses (Fig. 2B). Finally, live, single, EGFP-positive cells were analysed by the previously described PulSA technique, plotting the pulse width (W) versus height (H) of the EGFP fluorescence signal [38]. Cells expressing EGFP, which rarely contain inclusions (Fig. 2C; *left*), were used to set the gate to identify cells with inclusions formed by Fluc^WT^-EGFP (Fig. 2C; *centre*) or Fluc^DM^-EGFP (Fig. 2C; *right*).

**Fig. 2.**
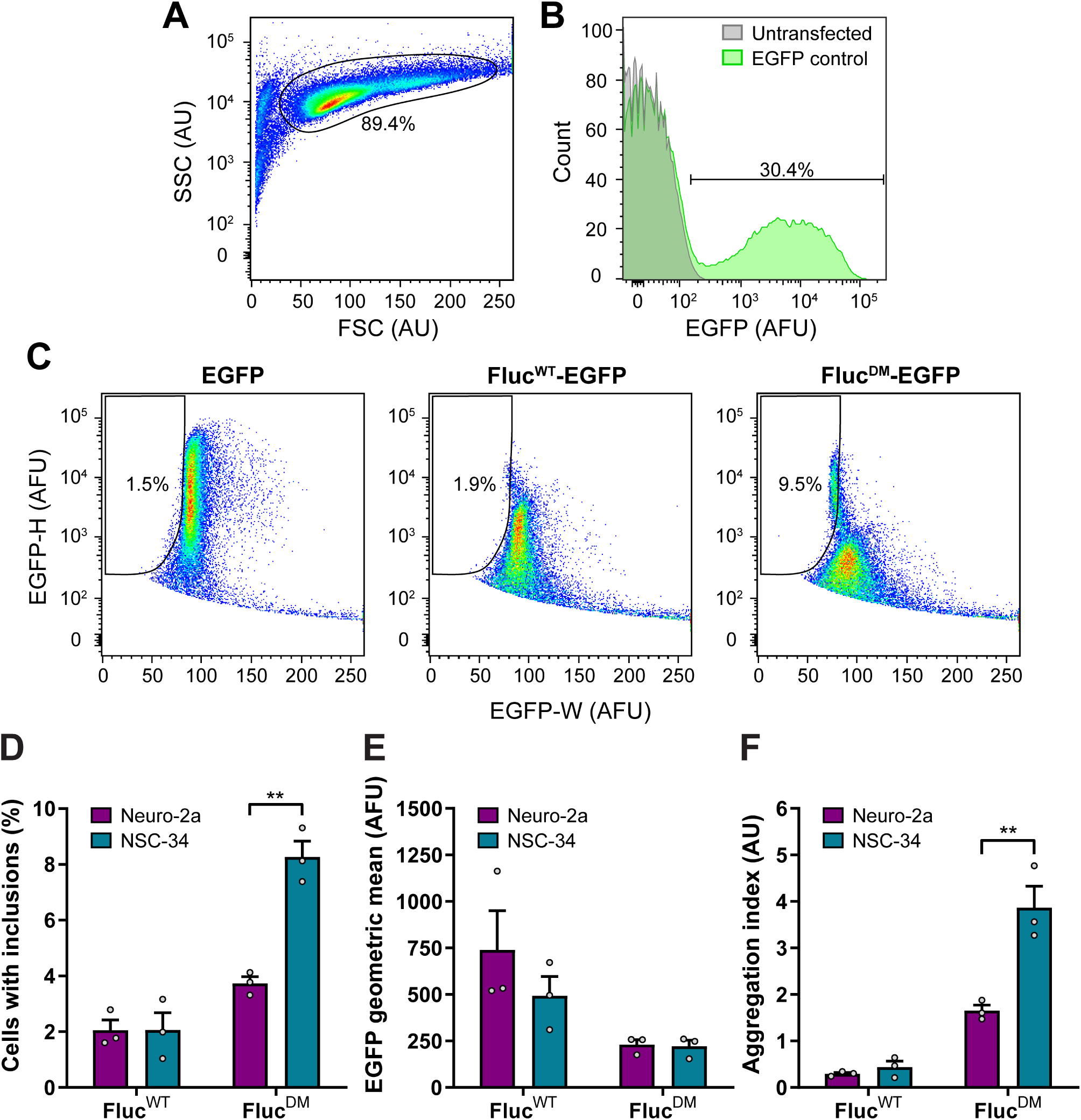
The formation of inclusions by Fluc in Neuro-2a and NSC-34 cells, as assessed by PulSA. Neuro-2a and NSC-34 cells were transiently transfected to express EGFP, Fluc^WT^-EGFP or Fluc^DM^-EGFP and were analysed by PulSA 48 h post-transfection. EGFP fluorescence was detected using a 488 nm laser and 525/50 nm emission filter and 100,000 events were collected for each cell population. Values within plots represent the percentage of cells within each gate. **(A)** Forward scatter (FSC) and side scatter (SSC) pseudocolour plot of untransfected cells (*left*). The polygonal gate encompasses the viable, live cells in the population which were selected for subsequent analyses. **(B)** Frequency histogram of the relative EGFP fluorescence of untransfected (*grey*) and transiently transfected cells expressing Fluc^DM^-EGFP (*green*). Cells expressing EGFP (gate shown) were selected for subsequent analyses. **(C)** Pulse shape analysis (PulSA) of cells transiently transfected with EGFP (*left*), Fluc^WT^-EGFP (*centre*) or Fluc^DM^-EGFP (*right*) used to identify cells with inclusions based upon the EGFP pulse width (W) and pulse height (H) signals. **(D)** The proportion of live, transfected cells with inclusions. **(E)** The geometric mean of EGFP, which corresponds to the amount of Fluc expressed within each cell population. **(F)** PulSA aggregation index, calculated as the ratio of the percentage of cells containing inclusions, divided by the geometric mean of EGFP fluorescence in EGFP-positive cells. Data in D – F is presented as the mean ± S.E.M (n=3 biological replicates of at least 10,000 cells). Significant differences between group means in the data were determined using a two-tailed Student’s t-test (\*\**P* < 0.01)

In Neuro-2a cells expressing Fluc^WT^-EGFP or Fluc^DM^-EGFP, the proportion of cells with inclusions was identified to be 2.0 ± 0.4% and 3.7 ± 0.2% respectively, whilst for NSC-34 cells the proportion of cells containing inclusions was 2.1 ± 0.6% and 8.3 ± 0.8% (Fig. 2D). To further compare the capacity of Neuro-2a and NSC-34 cells to prevent the aggregation of Fluc, an aggregation index was calculated. This aggregation index enabled the susceptibility of a particular cell type to the formation of inclusions by Fluc to be quantified. To account for the amount of Fluc expressed in the different cell populations, the geometric mean of the EGFP-positive cells was determined (Fig. 2E). There was no difference in the amount of Fluc^WT^ or Fluc^DM^ expressed between cell types. The PulSA aggregation index was calculated as the ratio of the percentage of cells identified to contain inclusions (as determined by PulSA) to the EGFP geometric mean of transfected cells. Based on these analyses, NSC-34 cells expressing Fluc^DM^ have a significantly higher aggregation index than Neuro-2a cells expressing Fluc^DM^ suggesting that NSC-34 cells have less capacity to prevent the aggregation of Fluc^DM^ into inclusions (Fig. 2F).

We next considered the relationship between inclusion formation and the amount of protein in individual cells as a complementary measure of the capacity of Neuro-2a and NSC-34 cells to suppress Fluc^DM^-EGFP aggregation. The flow cytometry data were binned based on the EGFP fluorescence intensity, grouping cells from each line with the same relative EGFP expression. Overall, 16 log-scale bins of equal size were used (Fig. 3A), and the proportion of cells with inclusions was then determined by PulSA in each bin independently. In all samples, cells in bins 1–10 did not contain inclusions. From bin 11 onwards, as the amount of Fluc expressed in cells increased there was a corresponding increase in the proportion of cells with inclusions (Fig. 3B). Neuro-2a cells expressing Fluc^DM^-EGFP were found to be less susceptible to inclusion formation by Fluc^DM^ than NSC-34 cells. This is observed as a significant shift to the right when bin number (e.g. bins 12 and 13) is plotted against the proportion of cells with inclusions. There was little difference in the proportion of cells containing inclusions in both cell lines when expressing Fluc^WT^, which remained below 40% even at the highest level of Fluc^WT^-EGFP expression (i.e. bins 14 – 16).

**Fig. 3.**
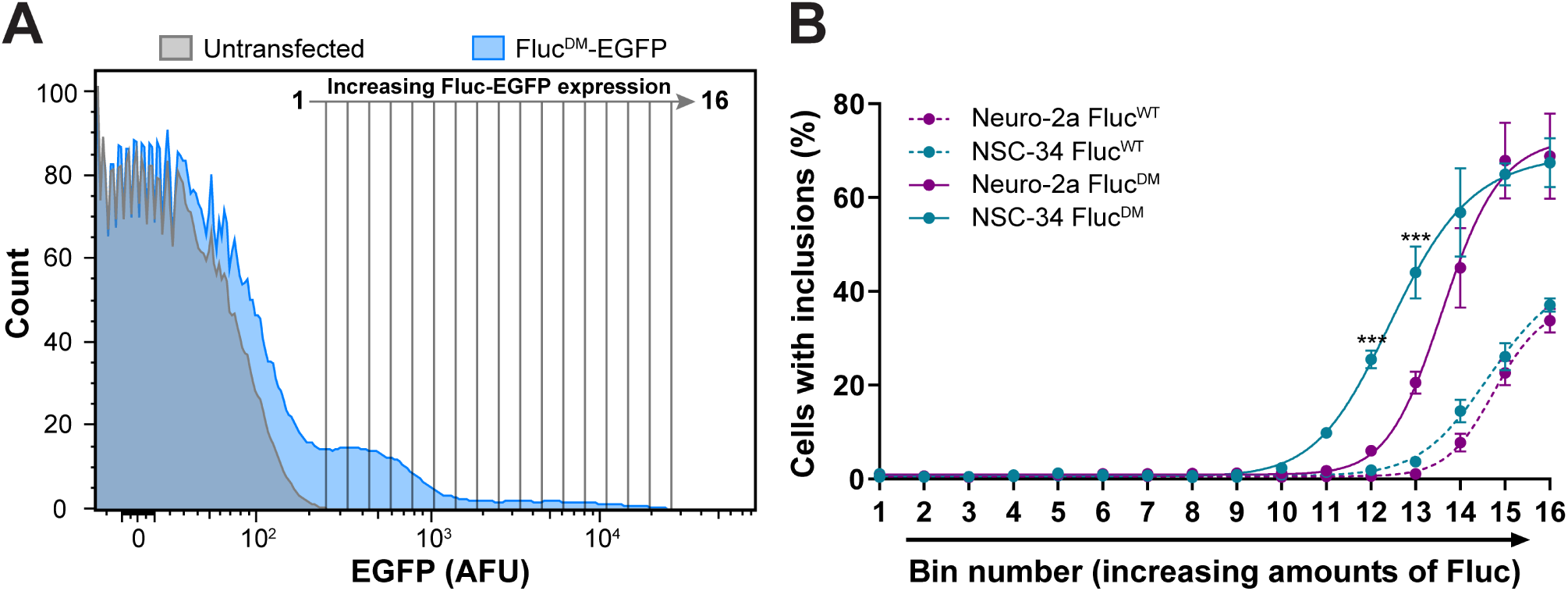
The relative susceptibility of Neuro-2a and NSC-34 cells to the formation of Fluc-based inclusions. Neuro-2a and NSC-34 cells were transiently transfected to express Fluc^WT^-EGFP or Fluc^DM^-EGFP and analysed by PulSA 48 h post-transfection. **(A)** Gating strategy used to determine the fraction of live cells with inclusions as a function of Fluc-EGFP expression. Overlay histograms of untransfected (*grey*) and Fluc^DM^-EGFP (*blue*) transfected cells were generated to identify transfected cells for subsequent analyses. The frequency histogram of EGFP fluorescence was then subdivided into 16 bins of equal size (based on a log scale), whereby bin 1 represents the lowest, and bin 16 the highest level of Fluc-EGFP expression. PulSA was then applied to obtain **(B)** the proportion of cells with inclusions in each bin. Data are presented as the mean ± S.E.M. of cells in each bin containing inclusions (n=3 biological replicates). Significant differences between group means were determined using a two-way ANOVA (*P* < 0.05) followed by a Bonferroni’s post-hoc test. Sample group means for cells expressing the same isoform of Fluc determined to be statistically different from each other within the same bin number are indicated (\*\*\**P* < 0.001)

### Proteomic analysis of Fluc^DM^ inclusions reveals cell-type specific engagement of the proteostasis machinery with aggregated proteins

We next sought to determine the molecular driver(s) of the susceptibility of NSC-34 cells to Fluc aggregation by examining the composition of Fluc^DM^ inclusions in each cell line. Neuro-2a and NSC-34 cells expressing Fluc^DM^-EGFP were harvested, lysed, and the insoluble proteins separated via gel electrophoresis. As a result of their large molecular mass, Fluc^DM^ inclusions migrate poorly and become trapped in the stacking gel during SDS-PAGE [44]. Consistent with this, cell lysates from Neuro-2a and NSC-34 cells expressing Fluc^DM^ contained a fluorescent band in the well of the stacking gel (Fig. 4A). The fluorescence intensity detected in the wells of the Neuro-2a samples was higher than that of the NSC-34 cells, which is likely due to differences in transfection efficiency between the two cells lines. Very low levels of fluorescence were also observed in the stacking gel of the untransfected cell lysates, likely due to autofluorescence from proteins such as collagen or elastin, cellular organelles such as mitochondria or lysosomes, or cyclic ring containing molecules, including NADPH or aromatic amino acids [45–47]. There were no discernible differences between cell types expressing Fluc^DM^ with regard to the profile of proteins that migrated within the resolving gel following SDS-PAGE (Fig. 4B). Proteins trapped in the stacking gel for both untransfected and Fluc-expressing cells were isolated for mass spectrometry-based quantitative proteomic analysis (regions indicated by the arrow).

**Fig. 4.**
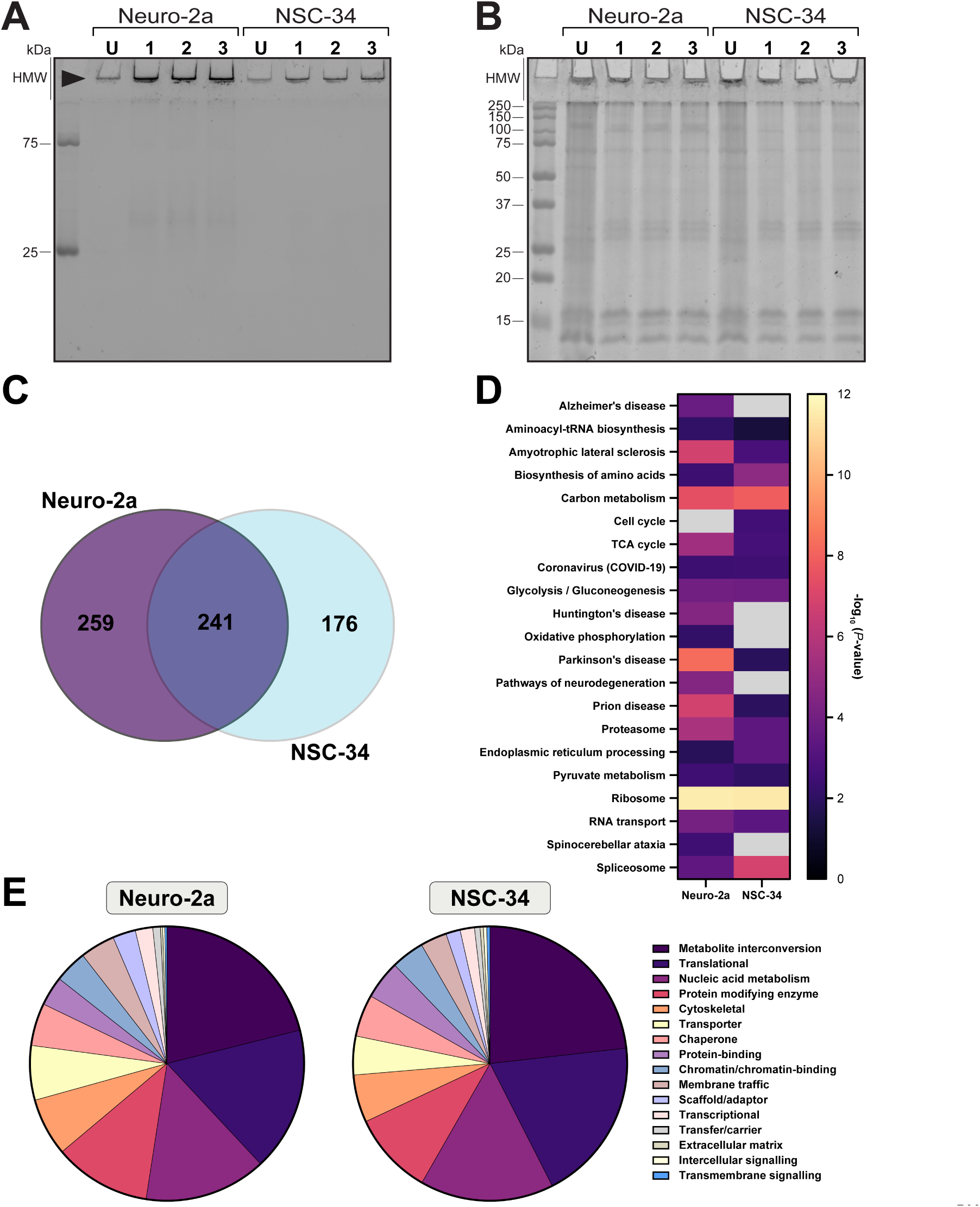
Distribution and enrichment of the KEGG pathways and classes of proteins identified within Fluc^DM^ inclusions by proteomic mass spectrometry. Neuro-2a and NSC-34 cells were transfected (or not) to express Fluc^DM^-EGFP and 48 h post-transfection were harvested, fractionated and the insoluble protein from each sample was separated by SDS-PAGE. **(A)** SDS-PAGE gel imaged following exposure to 488 nm light to detect fluorescence from EGFP-tagged Fluc^DM^. High molecular weight (HMW) proteins which are excited at 488 nm and are trapped in the stacking gel are indicated by the arrow. **(B)** SDS-PAGE gel stained with Coomassie Blue showing total protein in the insoluble lysates from these cells. Biological triplicates of cells transfected with Fluc^DM^-EGFP are shown, along with insoluble protein from cell lysates of untransfected Neuro-2a and NSC-34 cells (U). Molecular masses of the protein standards are shown to the left of the gel (in kDa). Proteins associated with Fluc^DM^ inclusions from Neuro-2a and NSC-34 cells were then identified by mass spectrometry. **(C)** The total number of proteins identified within Fluc^DM^ inclusions from Neuro-2a (*purple*) and NSC-34 (*blue*) cells. **(D)** The functional enrichment of KEGG pathways represented by proteins identified within Fluc^DM^ inclusions in Neuro-2a and NSC-34 cells. Only KEGG pathways that were significantly enriched are shown (threshold set to *P* ≤ 0.05). Colours represent –log_10_(*P*-values), whereby the highest values (highest enrichment) are in yellow, and the lowest values (lowest enrichment) are in black and greyed-out boxes represent a pathway not enriched in that cell type. **(E)** PANTHER analysis of protein classes most enriched within Fluc^DM^ inclusions in Neuro-2a (*left*) and NSC-34 (*right*) cells

Of the 676 proteins identified, 241 (35.7%) were detected in both cell lines, 259 (38.3%) were only detected in Neuro-2a cells and 176 (26%) were only detected in NSC-34 cells (Fig. 4C) (see Data Availability to access the full list of proteins identified). Proteins implicated in neurodegenerative diseases, such as Alzheimer’s disease, Parkinson’s disease, Huntington’s disease, spinocerebellar ataxia, ALS and prion-related diseases, were found to be enriched within the Fluc^DM^ inclusions in both Neuro-2a and NSC-34 cells (Fig. 4D). Other highly enriched KEGG pathways included those involved in transcription, translation, cell cycle control, respiration and metabolism, as well as major arms of the proteostasis network such as proteasomal degradation and protein processing in the ER. Consistent with this, the most highly represented protein classes in both Neuro-2a and NSC-34 cells were metabolic enzymes, RNA binding proteins (i.e. translational machinery), cytoskeletal proteins, proteins involved in transport and chaperones (PANTHERdb; Fig. 4E). Overall, the distribution of protein classes found to be associated with Fluc^DM^ inclusions were nearly identical between the cell lines.

We next examined the interaction networks for proteins identified within Fluc^DM^ inclusions unique to, or common between, Neuro-2a and NSC-34 cells. Proteins were manually clustered based on protein class; the major classes of proteins identified were similar even among proteins unique to Neuro-2a (Fig. 5A) and NSC-34 (Fig. 5B) cells. The central hubs consisted of proteins involved in RNA metabolism, translation, protein modification, metabolite interconversion, cytoskeletal organisation, proteasomal degradation and chaperone proteins; categories which were also represented among the proteins common to both cell lines (Fig. 5C). Interestingly, proteins involved in ER processing were only identified in aggregates derived from Neuro-2a cells (Fig. 5A). Analyses confirmed that the protein-protein interaction enrichment was statistically significant across the three networks (i.e. *P* < 1.0 × 10^-16^, in all cases). This indicates that these proteins have more interactions among themselves than would be expected by chance compared to a random set of proteins from the genome with the same size and distribution. Despite this broad similarity in protein classes associated with aggregation, the specific proteins identified in aggregates from each cell line varied, suggesting differences in the specific mechanisms by which these cell lines respond to Fluc^DM^ aggregation. The relative abundance of proteins identified in inclusions common to both cell lines was similarly heterogeneous (Fig. 5C). The most highly represented protein classes were ribosomal proteins, translational machinery, metabolite interconversion enzymes, proteasome components and chaperone proteins. These protein classes, all of which are components of the proteostasis network, act as interaction hubs, linking all the proteins found to associate with Fluc^DM^ inclusions in both cell lines.

**Fig. 5.**
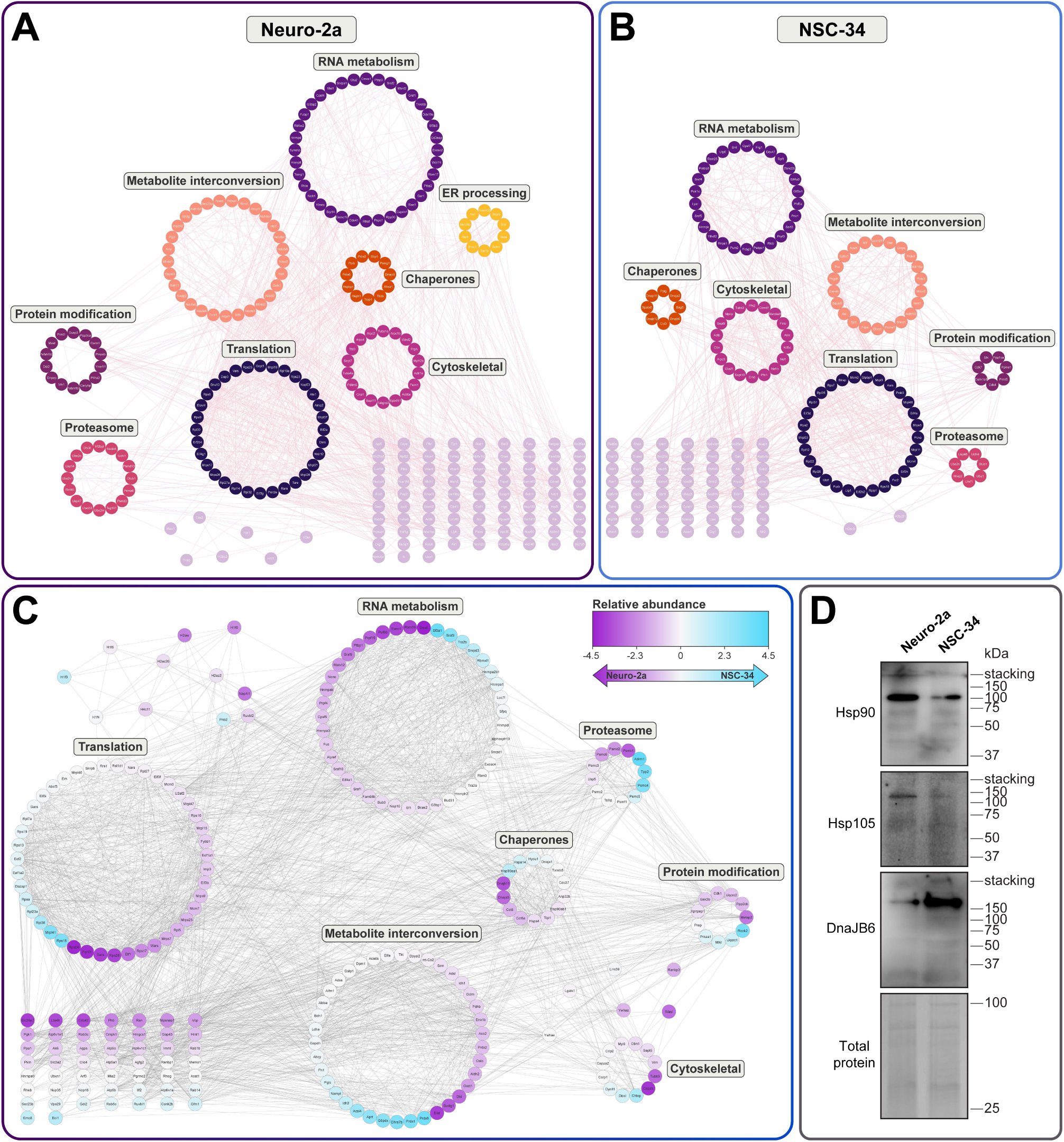
STRING analysis and relative abundance of proteins identified within Fluc^DM^ inclusions in either Neuro-2a or NSC-34 cells, or both cell lines. Protein-protein interaction networks for proteins identified in Fluc^DM^ inclusions from **(A)** Neuro-2a, **(B)** NSC-34 cells, or **(C)** both Neuro-2a and NSC-34 cells. Networks were constructed using the STRING application in Cytoscape. Nodes represent individual proteins manually clustered according to class, joined by edges indicating known or predicted protein-protein interactions (STRING score > 0.4). Nodes are coloured by (A, B) class or (C) the fold change in abundance (NSC-34/Neuro-2a), whereby nodes in purple represent proteins more highly expressed in Neuro-2a cells and nodes in blue represent proteins more highly expressed in NSC-34 cells. **(D)** Confirmation of proteins identified to associate with insoluble Fluc^DM^ in either Neuro-2a or NSC-34 cells, or both cell lines, by immunoblotting. Neuro-2a and NSC-34 cells were transiently transfected to express Fluc^DM^-EGFP and analysed 48 h post-transfection by NP-40 fractionation and subsequent immunoblotting. In the insoluble fraction, antibodies against Hsp90, Hsp105 and DnaJB6 were used to detect endogenous expression of the corresponding Hsps. Coomassie-stained total protein was used as a loading control. Molecular masses of the protein standards are shown to the right (in kDa)

Three proteins found in Fluc^DM^ inclusions were chosen for further validation; one identified in both cell lines (heat shock protein 90, Hsp90), one uniquely found in inclusions from Neuro-2a cells (heat shock protein 105, Hsp105), and one uniquely found in inclusions from NSC-34 cells (the heat shock protein 40 isoform, DnaJB6) (Fig. 5D). Immunoblotting confirmed that Hsp90 was detected in the insoluble fraction of both Neuro-2a and NSC-34 cells. Immunoblotting also confirmed the association of Hsp105 with Fluc^DM^ inclusions only in Neuro-2a cells. Finally, DnaJB6 was found at high levels in the insoluble cell lysate of NSC-34 cells at a high molecular weight (>250 kDa), with much lower levels detected in the insoluble cell lysate of Neuro-2a cells. This could suggest that DnaJB6 forms a complex with Fluc^DM^ in inclusions and that these interactions are not disrupted by the experimental conditions used in this work. Overall, the results of the immunoblotting validate the presence of those proteins that were identified to co-aggregate with Fluc^DM^ by mass spectrometry.

### Gene ontology analysis of Neuro-2a and NSC-34 proteomes reveals differential expression of ER unfolded protein response components within the proteostasis network

To determine whether there are differences in the proteins expressed between Neuro-2a and NSC-34 cells across various arms of the proteostasis network, gene ontology mapping was undertaken using previously published data on the whole proteomes of these two cell types [37]. Analysis of the entire 4,325 proteins (Fig. 6A (i)) identified in Neuro-2a and NSC-34 cells revealed a strong positive linear correlation between the abundance of an individual protein in each cell line (Fig. 6A (ii)). An alternate way to measure this relationship is via the ratio of protein abundance in Neuro-2a compared to NSC-34 cells (Fig. 6A (iii)). That the log_2_ transformation of this ratio was centred on 0 is consistent with no significant difference in the expression of the proteome between the two cells lines. We hypothesised that any arms of the proteostasis network which deviate from this relationship may contribute to differences in the aggregation susceptibility of each cell line. To explore this, we first collected subsets of the proteome categorised according to their association with gene ontology terms relevant to specific arms of the proteostasis network. For example, there were 74 proteins associated with the term “chaperone” present in the dataset, the ubiquitin-proteosome system, represented by the terms “ubiquitin” or “proteasome”, was associated with 441 proteins, while the term “autophagy” was associated with 115 proteins (Fig. 6B-D (i)). Evaluation of these arms of the proteostasis network indicated that there was no significant difference in the correlation (Fig. 6B-D (ii)) or distribution of protein abundance (Fig. 6B-D (iii)) between Neuro-2a and NSC-34 cells.

**Fig. 6.**
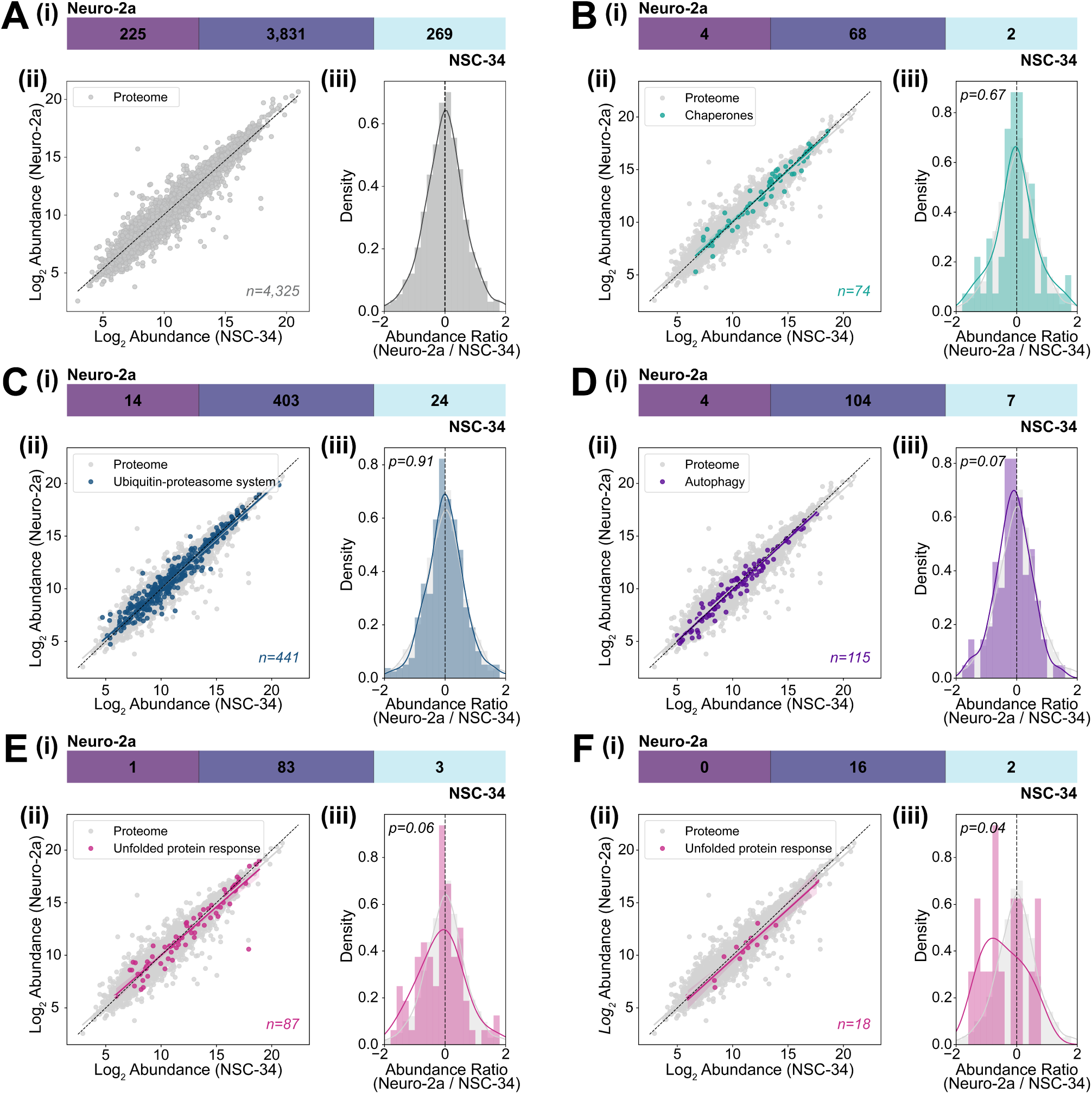
Gene ontology mapping of proteostasis pathways in Neuro-2a and NSC-34 cells compared to the whole proteome. **(A)** The relationship between protein abundance in Neuro-2a and NSC-34 cells [37] was examined. Proteins were also mapped to central proteostasis hubs using gene ontology terms. Ontology terms were selected based on their inclusion of the keywords **(B) “**chaperones”, **(C)** “ubiquitin” “proteasome” system, **(D)** “autophagy”, **(E) “**unfolded protein” response, or the specific term **(F) “**GO:0030968 (ER unfolded protein response)”. Panel (i) shows the total number of proteins identified in the proteome or following gene ontology mapping within Neuro-2a (*purple*) and NSC-34 (*blue*) cells. Panel (ii) shows the log_2_ abundance of all proteins quantified in Neuro-2a and NSC-34 cells (*grey*), optionally overlayed with the proteins corresponding to each hub term (B – F; *coloured*), fitted with linear regression. *n* reports the number of proteins in the hub term. Panel (iii) shows the distribution of abundance ratios (log_2_[Neuro-2a/NSC-34]) for the whole proteome (*grey*), optionally overlayed with the proteins corresponding to individual hub terms (B – F; *coloured*). Distributions were analysed by Wilcoxon test, with *P* ≤ 0.05 considered statistically significant

The final category we considered was the “unfolded protein response”, associated with 87 proteins in the dataset (Fig. 6E (i)). The abundance of proteins identified within this pathway was highly correlated between Neuro-2a and NSC-34 cells; however, the slope of the correlation deviated from that of the whole proteome (Fig. 6E (ii)). The distribution of ratios for the unfolded protein response compared to the proteome also appeared left skewed despite this difference not reaching statistical significance (*P* = 0.06) (Fig. 6E (iii)). This prompted us to examine individual ontology terms relevant to the unfolded protein response, including GO:0030968 (ER unfolded protein response) associated with 18 proteins expressed in Neuro-2a and NSC-34 cells (Fig. 6F (i)). The abundance of these proteins between Neuro-2a and NSC-34 cells was highly correlated, similar to that of the whole proteome (Fig. 6F (ii)). However, the distribution of the expression ratio was significantly left-skewed compared to the whole proteome (*P* = 0.04; Fig. 6F (iii)). Given the relatively small sample size, bootstrapping was performed in which the equivalent size was randomly selected from the proteome 1,000 times. In only 43 examples was the *P*-value as low (i.e. < 0.04) as the present case, corresponding to a less than 5% chance of the values between Neuro-2a and NSC-34 cells being this different in a random collection of proteins from the proteome. Together, these results indicate that the ER unfolded protein response in Neuro-2a and NSC-34 cells is differentially expressed between these two cell lines. This may account for the difference in the propensity of Fluc^DM^ to form inclusions between these two cell lines.

## Discussion

As the accumulation of aggregation-prone proteins into inclusions is a pathological hallmark of many diseases, it is critical to understand the precise roles various components of the proteostasis network have in preventing this process. Moreover, quantitatively assessing the capacity of a cell to prevent an aggregation-prone protein from forming inclusions remains one of the major challenges in the field. The ability to quantitatively measure the proteostasis capacity of a cell is an important step towards deciphering why some cell types are more susceptible to the formation of inclusions (and cause disease) than others. Furthermore, the deterioration of proteostasis capacity with age is believed to be related to an increase in toxic protein aggregation that is associated with disease [34, 35].

In this work, flow cytometric analyses of cells expressing aggregation-prone protein Fluc^DM^ demonstrates that NSC-34 cells are more susceptible to inclusion formation by Fluc^DM^ than Neuro-2a cells. Thus, the results demonstrate that lower amounts of Fluc^DM^ are required before inclusions start to form in NSC-34 cells compared to Neuro-2a cells. The level of expression of an aggregation-prone protein is known to impact the formation of inclusions, with proteins forming inclusions more readily when expressed at high levels [38, 48–51]. By considering the level of Fluc^DM^ expressed in the cell population, the aggregation index derived from this work enables comparison between different cell types.

One potential reason NSC-34 cells are more susceptible to cytotoxic inclusion formation than Neuro-2a cells is that they are less equipped at the proteome level to deal with aggregation-prone proteins. Previous work has indicated that the global proteomes of Neuro-2a and NSC-34 cells are very similar [37]; however, the types of proteins that readily engage with an aggregation-prone protein as it forms inclusions in these cells has never been compared. Differences in the classes of proteins found within proteinaceous deposits may explain why some cell types can more readily prevent inclusion formation [52]. Moreover, the accumulation of proteins into inclusions can lead to the functional loss of key components of the proteostasis network. Mass spectrometry analysis revealed that the classes of proteins trapped within Fluc^DM^ inclusions in Neuro-2a and NSC-34 cells are near identical. These protein classes include important components of the systems that maintain intracellular proteostasis, including proteins related to processing in the ER, chaperones and proteasomal machinery.

The most enriched classes of proteins identified in Fluc^DM^ inclusions in both Neuro-2a and NSC-34 cells were those associated with the translational machinery. Indeed, previous work has suggested that disruption of translation is associated with neurodegenerative disorders. For example, Halliday [53] identified a link between the expression of misfolded forms of prion and tau proteins, and translational repression *in vivo*. Translation-relevant terms were also reported by Kamelgarn [54] in a gene ontology assessment of the proteins contained within cytoplasmic inclusions formed by the ALS– and frontotemporal dementia-associated fused in sarcoma (FUS) protein. Moreover, in a humanised mouse model of ALS/frontotemporal dementia, the expression of mutant FUS inhibited protein synthesis and led to impaired neuronal synaptic function [55]. Furthermore, the unfolded protein response pathway, which mediates the rate of protein synthesis in cells, is over-activated in the brains of patients with neurodegenerative diseases [53, 55–58] and therefore may be a promising target for pharmacological intervention to alleviate translational repression in neurodegeneration. The identification of translational machinery within Fluc^DM^ inclusions reinforces the notion that the aggregation of proteins into inclusions disrupts protein translation and this may be associated with neurodegenerative disease pathogenesis.

Another highly represented class of proteins found within Fluc^DM^ inclusions in both cell lines were chaperones, including members of the Hsp40, Hsp70, Hsp90, Hsp110 families and the chaperonin TriC/CCT. Previous work has demonstrated the presence of chaperone proteins localised within proteinaceous deposits, potentially due to their roles in maintaining the solubility and function of aggregation-prone proteins. For example, Hsp70 and Hsp40 were found to be associated with inclusions formed by polyQ-expanded ataxin-1 [59], ataxin-3 [60] and huntingtin [61]. Intracellular inclusions formed by mutant superoxide dismutase 1 (SOD1) contain αB-crystallin and Hsc70 (i.e. constitutively expressed Hsp70) [52]. Hsp70 has been identified in protein aggregates extracted from the brains of ALS/frontotemporal dementia patients with transactive response DNA-binding protein 43 (TDP-43)-positive proteinopathies [62]. Furthermore, Hsp27 and αB-crystallin were detected within neurofibrillary tangles of patients with Alzheimer’s disease [63]. Chaperone proteins are likely found within inclusions due to their roles in maintaining the solubility and function of aggregation-prone proteins. Thus, the presence of chaperones in inclusions may be due to them having an important role in mitigating the toxicity of inclusions (e.g. through disaggregation) or may simply reflect a failure to prevent protein aggregation.

A variety of proteins that play a role in proteostasis were found to be associated with Fluc^DM^ inclusions in both Neuro-2a and NSC-34 cells. These included proteasome components and chaperones. Thus, whilst there were few differences between Neuro-2a and NSC-34 cells with regard to the classes of proteins that associate with Fluc^DM^ in inclusions formed in these cells, these data highlight the important role that protein quality control pathways of the proteostasis network play in attempting to mitigate cytotoxic protein aggregation.

Interestingly, the STRING analysis determined that the clustering of proteins in Neuro-2a and NSC-34 cells were largely similar, with the exception of proteins related to ER processing, a pathway which was not enriched in NSC-34 cells. This prompted further investigation into the individual protein quality control pathways in Neuro-2a and NSC-34 cells, and in particular, the ER unfolded protein response.

Gene ontology mapping of proteins associated with the ER unfolded protein response expressed in Neuro-2a and NSC-34 cells determined that these proteins are regulated differently when compared to the whole proteome. Taken together, these data indicate that proteins related to ER processing do not associate with Fluc^DM^ in inclusions in NSC-34 cells as frequently compared to Neuro-2a cells. This may help to explain the susceptibility of NSC-34 cells to inclusion formation, when compared to Neuro-2a cells. Moreover, the significant difference in the distribution of proteins associated with this pathway, when compared to the whole proteome, may signify a generalised dysfunction of the ER unfolded protein response in neurons. The ER is an important component of the cellular quality control network that helps to ensure correct folding of many newly synthesised secretory and transmembrane proteins. Perturbations in ER homeostasis can result in an accumulation of unfolded proteins in the ER lumen, termed ER stress [64], which triggers a dynamic signalling pathway known as the unfolded protein response [65]. Ongoing ER stress, owing to mutations in disease-related genes or malfunctions in the secretory pathway, can lead to cell death and, in some cases, neurodegeneration [66, 67].

Overall, this work successfully exploited the aggregation-prone nature of Fluc^DM^ to quantitatively ascertain that NSC-34 cells are more susceptible to inclusion formation by destabilised proteins than Neuro-2a cells. Moreover, the methods presented here provide a framework towards comparing the proteostasis capacity of many cell types. The proteins identified within Fluc^DM^-containing proteinaceous deposits in Neuro-2a and NSC-34 cells were near identical, with the major classes belonging to the protein quality control network, highlighting the important role these pathways play in maintaining the solubility of the protein in cells. Identification of the pathways important in preventing protein aggregation, along with techniques able to assess the susceptibility of cell types to inclusion formation, provides a basis for the development of effective therapies against neurodegenerative diseases

## Acknowledgements

We thank staff at Molecular Horizons for technical and administrative support.

## Author contributions

**Shannon McMahon:** Methodology, Formal analysis, Investigation, Data Curation, Writing – Original Draft, Visualization. **Dezerae Cox:** Software, Formal analysis, Data Curation, Writing – Review & Editing. **Flora Cheng:** Methodology, Investigation, Writing – Review & Editing. **Albert Lee:** Methodology, Formal analysis. Resources, Writing – Review & Editing. **Justin J Yerbury:** Conceptualization, Supervision. **Heath Ecroyd:** Conceptualization, Writing – Review & Editing, Supervision, Project administration, Funding acquisition.

## Data availability

The mass spectrometry proteomics data have been deposited to the ProteomeXchange via the PRIDE partner repository with the identifier PXD031826 (PubMed ID: 34723319).

## Dedication

The authors would like to dedicate the work presented in this manuscript to their dear friend and colleague, the late Professor Justin J Yerbury.

## References

[1] Hartl FU, Hayer-Hartl M (2002) Molecular chaperones in the cytosol: from nascent chain to folded protein. Science 295:1852–1858. 10.1126/science.1068408

[2] Wang W, Nema S, Teagarden D (2010) Protein aggregation – pathways and influencing factors. Int J Pharm 390:89–99. 10.1016/j.ijpharm.2010.02.025

[3] Ecroyd H, Carver JA (2008) Unraveling the mysteries of protein folding and misfolding. IUBMB Life 60:769–774. 10.1002/iub.117

[4] Fink AL (1998) Protein aggregation: folding aggregates, inclusion bodies and amyloid. Folding and Design 3:R9–R23. 10.1016/S1359-0278(98)00002-9

[5] Stefani M (2008) Protein folding and misfolding on surfaces. International Journal of Molecular Sciences 9:2515–2542. 10.3390/ijms9122515

[6] Shea JE, Brooks CL (2001) From folding theories to folding proteins: a review and assessment of simulation studies of protein folding and unfolding. Annual Review of Physical Chemistry 52:499–535. 10.1146/annurev.physchem.52.1.499

[7] Dobson CM (2004) Experimental investigation of protein folding and misfolding. Methods 34:4–14. 10.1016/j.ymeth.2004.03.002

[8] Stranks SD, Ecroyd H, Van Sluyter S, Waters EJ, Carver JA, Von Smekal L (2009) Model for amorphous aggregation processes. Physical Review E 80:1–13. 10.1103/PhysRevE.80.051907

[9] Greenwald J, Riek R (2010) Biology of amyloid: structure, function, and regulation. Structure 18:1244–1260. 10.1016/j.str.2010.08.009

[10] Wigley CW, Fabunmi RP, Lee MG, Marino CR, Muallem S, DeMartino GN, Thomas PJ (1999) Dynamic association of proteasomal machinery with the centrosome. J Cell Biol 145:481–490. 10.1083/jcb.145.3.481

[11] Kirstein J, Strahl H, Molière N, Hamoen LW, Turgay K (2008) Localization of general and regulatory proteolysis in Bacillus subtilis cells. Mol Microbiol 70:682–694. 10.1111/j.1365-2958.2008.06438.x

[12] Kaganovich D, Kopito R, Frydman J (2008) Misfolded proteins partition between two distinct quality control compartments. Nature 454:1088–1095. 10.1038/nature07195

[13] Cecchi C, Baglioni S, Fiorillo C, Pensalfini A, Liguri G, Nosi D, Rigacci S, Bucciantini M, Stefani M (2005) Insights into the molecular basis of the differing susceptibility of varying cell types to the toxicity of amyloid aggregates. J Cell Sci 118:3459–3470. 10.1242/jcs.02473

[14] Wyttenbach A, Sauvageot O, Carmichael J, Diaz-Latoud C, Arrigo AP, Rubinsztein DC (2002) Heat shock protein 27 prevents cellular polyglutamine toxicity and suppresses the increase of reactive oxygen species caused by huntingtin. Hum Mol Genet 11:1137–1151.

[15] Kundra R, Dobson CM, Vendruscolo M (2020) A cell– and tissue-specific weakness of the protein homeostasis system underlies brain vulnerability to protein aggregation. iScience 23:100934. 10.1016/j.isci.2020.100934

[16] Chen YR, Harel I, Singh PP, Ziv I, Moses E, Goshtchevsky U, Machado BE, Brunet A, Jarosz DF (2024) Tissue-specific landscape of protein aggregation and quality control in an aging vertebrate. Dev Cell 59:1892–1911.e13. 10.1016/j.devcel.2024.04.014

[17] Park JH, Nordström U, Tsiakas K, Keskin I, Elpers C, Mannil M, Heller R, Nolan M, Alburaiky S, Zetterström P, Hempel M, Schara-Schmidt U, Biskup S, Steinacker P, Otto M, Weishaupt J, Hahn A, Santer R, Marquardt T, Marklund SL, Andersen PM (2023) The motor system is exceptionally vulnerable to absence of the ubiquitously expressed superoxide dismutase-1. Brain Communications 5. 10.1093/braincomms/fcad017

[18] Lee SH, Du J, Hwa J, Kim WH (2020) Parkin coordinates platelet stress response in diabetes mellitus: A big role in a small cell. International Journal of Molecular Sciences 21:1–19. 10.3390/ijms21165869

[19] Lim J, Yue Z (2015) Neuronal aggregates: formation, clearance, and spreading. Dev Cell 32:491–501. 10.1016/j.devcel.2015.02.002

[20] Saxena S, Caroni P (2011) Selective neuronal vulnerability in neurodegenerative diseases: from stressor thresholds to degeneration. Neuron 71:35–48. 10.1016/j.neuron.2011.06.031

[21] Smith HL, Li W, Cheetham ME (2015) Molecular chaperones and neuronal proteostasis. Seminars in Cell and Developmental Biology 40:142–152. 10.1016/j.semcdb.2015.03.003

[22] Gebert N, Rahman S, Lewis CA, Ori A, Cheng CW (2021) Identifying cell-type-specific metabolic signatures using transcriptome and proteome analyses. Current Protocols 1:e245. 10.1002/cpz1.245

[23] Kampinga HH, Bergink S (2016) Heat shock proteins as potential targets for protective strategies in neurodegeneration. The Lancet Neurology 14:749–759. 10.1016/S1474-4422(16)00099-5

[24] Malhotra JD, Kaufman RJ (2007) The endoplasmic reticulum and the unfolded protein response. Seminars in Cell and Developmental Biology 18:716–731. 10.1016/j.semcdb.2007.09.003

[25] Rutkowski DT, Kaufman RJ (2007) That which does not kill me makes me stronger: adapting to chronic ER stress. Trends Biochem Sci 32:469–476. 10.1016/j.tibs.2007.09.003

[26] Matus S, Glimcher LH, Hetz C (2011) Protein folding stress in neurodegenerative diseases: a glimpse into the ER. Curr Opin Cell Biol 23:239–252. 10.1016/j.ceb.2011.01.003

[27] Morimoto RI (2008) Proteotoxic stress and inducible chaperone networks in neurodegenerative disease and aging. Genes and Development 22:1427–1438. 10.1101/gad.1657108

[28] Finkbeiner S, Cuervo AM, Morimoto RI, Muchowski PJ (2006) Disease-modifying pathways in neurodegeneration. J Neurosci 26:10349–10357. 10.1523/JNEUROSCI.3829-06.2006

[29] Komatsu M, Waguri S, Chiba T, Murata S, Iwata JI, Tanida I, Ueno T, Koike M, Uchiyama Y, Kominami E, Tanaka K (2006) Loss of autophagy in the central nervous system causes neurodegeneration in mice. Nature 441:880–884. 10.1038/nature04723

[30] Gidalevitz T, Ben-Zvi A, Ho KH, Brignull HR, Morimoto RI (2006) Progressive disruption of cellular protein folding in models of polyglutamine diseases. Science 311:1471–1474. 10.1126/science.1124514

[31] Hutt DM, Powers ET, Balch WE (2009) The proteostasis boundary in misfolding diseases of membrane traffic. FEBS Lett 583:2639–2646. 10.1016/j.febslet.2009.07.014

[32] Powers ET, Morimoto RI, Dillin A, Kelly JW, Balch WE (2009) Biological and chemical approaches to diseases of proteostasis deficiency. Annu Rev Biochem 78:959–991. 10.1146/annurev.biochem.052308.114844

[33] Gidalevitz T, Kikis EA, Morimoto RI (2010) A cellular perspective on conformational disease: the role of genetic background and proteostasis networks. Curr Opin Struct Biol 20:23–32. 10.1016/j.sbi.2009.11.001

[34] Brehme M, Voisine C, Rolland T, Wachi S, Soper JH, Zhu Y, Orton K, Villella A, Garza D, Vidal M, Ge H, Morimoto RI (2014) A chaperome subnetwork safeguards proteostasis in aging and neurodegenerative disease. Cell Reports 9:1135–1150. 10.1016/j.celrep.2014.09.042

[35] Hipp MS, Park SH, Hartl FU (2014) Proteostasis impairment in protein-misfolding and –aggregation diseases. Trends Cell Biol 24:506–514. 10.1016/j.tcb.2014.05.003

[36] Gupta R, Kasturi P, Bracher A, Loew C, Zheng M, Villella A, Garza D, Hartl FU, Raychaudhuri S (2011) Firefly luciferase mutants as sensors of proteome stress. Nat Methods 8:879–884. 10.1038/nmeth.1697

[37] Hornburg D, Drepper C, Butter F, Meissner F, Sendtner M, Mann M (2014) Deep proteomic evaluation of primary and cell line motoneuron disease models delineates major differences in neuronal characteristics. Mol Cell Proteomics 13:3410–3420. 10.1074/mcp.M113.037291

[38] Ramdzan YM, Polling S, Chia CPZ, Ng IHW, Ormsby AR, Croft NP, Purcell AW, Bogoyevitch MA, Ng DCH, Gleeson PA, Hatters DM (2012) Tracking protein aggregation and mislocalization in cells with flow cytometry. Nat Methods 9:467–470. 10.1038/nmeth.1930

[39] Raudvere U, Kolberg L, Kuzmin I, Arak T, Adler P, Peterson H, Vilo J (2019) g: Profiler: a web server for functional enrichment analysis and conversions of gene lists. Nucleic Acids Res 47:W191–W198.

[40] Szklarczyk D, Morris JH, Cook H, Kuhn M, Wyder S, Simonovic M, Santos A, Doncheva NT, Roth A, Bork P, Jensen LJ, Von Mering C (2017) The STRING database in 2017: quality-controlled protein-protein association networks, made broadly accessible. Nucleic Acids Res 45:D362–D368. 10.1093/nar/gkw937

[41] Shannon P, Markiel A, Ozier O, Baliga NS, Wang JT, Ramage D, Amin N, Schwikowski B, Ideker T (2003) Cytoscape: a software environment for integrated models of biomolecular interaction networks. Genome Res 13:2498–2504. 10.1101/gr.1239303

[42] Croft D, Mundo AF, Haw R, Milacic M, Weiser J, Wu G, Caudy M, Garapati P, Gillespie M, Kamdar MR, Jassal B, Jupe S, Matthews L, May B, Palatnik S, Rothfels K, Shamovsky V, Song H, Williams M, Birney E, Hermjakob H, Stein L, D’Eustachio P (2014) The Reactome pathway knowledgebase. Nucleic Acids Res 42:D472–D477. 10.1093/nar/gkt1102

[43] Towbin H, Staehelin T, Gordon J (1979) Electrophoretic transfer of proteins from polyacrylamide gels to nitrocellulose sheets: procedure and some applications. Proceedings of the National Academy of Sciences of the United States of America 76:4350–4354.

[44] Shevchenko A, Tomas H, Havliš J, Olsen JV, Mann M (2007) In-gel digestion for mass spectrometric characterization of proteins and proteomes. Nature Protocols 1:2856–2860. 10.1038/nprot.2006.468

[45] Andersson H, Baechi T, Hoechl M, Richter C (1998) Autofluorescence of living cells. Journal of Microscopy 191:1–7. 10.1046/j.1365-2818.1998.00347.x

[46] Fujimoto D, Akiba Ky, Nakamura N (1977) Isolation and characterization of a fluorescent material in bovine achilles tendon collagen. Biochem Biophys Res Commun 76:1124–1129. 10.1016/0006-291X(77)90972-X

[47] Blomfield J, Farrar JF (1969) The fluorescent properties of maturing arterial elastin. Cardiovascular Research 3:161–170. 10.1093/cvr/3.2.161

[48] Ramdzan YM, Wood R, Hatters DM (2013) Pulse shape analysis (PulSA) to track protein translocalization in cells by flow cytometry: applications for polyglutamine aggregation. Methods in Molecular Biology 1017:85–93. 10.1007/978-1-62703-438-8_6

[49] Ormsby AR, Ramdzan YM, Mok YF, Jovanoski KD, Hatters DM (2013) A platform to view huntingtin exon 1 aggregation flux in the cell reveals divergent influences from chaperones hsp40 and hsp70. J Biol Chem 288:37192–37203. 10.1074/jbc.M113.486944

[50] Arrasate M, Mitra S, Schweitzer ES, Segal MR, Finkbeiner S (2004) Inclusion body formation reduces levels of mutant huntingtin and the risk of neuronal death. Nature 431:805–810. 10.1038/nature02998

[51] Ciryam P, Kundra R, Morimoto RI, Dobson CM, Vendruscolo M (2015) Supersaturation is a major driving force for protein aggregation in neurodegenerative diseases. Trends Pharmacol Sci 36:72–77. 10.1016/j.tips.2014.12.004

[52] Bergemalm D, Forsberg K, Srivastava V, Graffmo KS, Andersen PM, Brännström T, Wingsle G, Marklund SL (2010) Superoxide dismutase-1 and other proteins in inclusions from transgenic amyotrophic lateral sclerosis model mice. J Neurochem 114:408–418. 10.1111/j.1471-4159.2010.06753.x

[53] Halliday M, Radford H, Zents KAM, Molloy C, Moreno JA, Verity NC, Smith E, Ortori CA, Barrett DA, Bushell M, Mallucci GR (2017) Repurposed drugs targeting eIF2α-P-mediated translational repression prevent neurodegeneration in mice. Brain 140:1768–1783. 10.1093/brain/awx074

[54] Kamelgarn M, Chen J, Kuang L, Jin H, Kasarskis EJ, Zhu H (2018) ALS mutations of FUS suppress protein translation and disrupt the regulation of nonsense-mediated decay. Proceedings of the National Academy of Sciences of the United States of America 115:E11904–E11913. 10.1073/pnas.1810413115

[55] López-Erauskin J, Tadokoro T, Baughn MW, Myers B, McAlonis-Downes M, Chillon-Marinas C, Asiaban JN, Artates J, Bui AT, Vetto AP, Lee SK, Le AV, Sun Y, Jambeau M, Boubaker J, Swing D, Qiu J, Hicks GG, Ouyang Z, Fu XD, Tessarollo L, Ling SC, Parone PA, Shaw CE, Marsala M, Lagier-Tourenne C, Cleveland DW, Da Cruz S (2018) ALS/FTD-linked mutation in FUS suppresses intra-axonal protein synthesis and drives disease without nuclear loss-of-function of FUS. Neuron 100:816–830. 10.1016/j.neuron.2018.09.044

[56] Bosco DA (2018) Translation dysregulation in neurodegenerative disorders. Proceedings of the National Academy of Sciences of the United States of America 115:12842–12844. 10.1073/pnas.1818493115

[57] Zhang YJ, Gendron TF, Ebbert MTW, O’Raw AD, Yue M, Jansen-West K, Zhang X, Prudencio M, Chew J, Cook CN, Daughrity LM, Tong J, Song Y, Pickles SR, Castanedes-Casey M, Kurti A, Rademakers R, Oskarsson B, Dickson DW, Hu W, Gitler AD, Fryer JD, Petrucelli L (2018) Poly(GR) impairs protein translation and stress granule dynamics in C9orf72-associated frontotemporal dementia and amyotrophic lateral sclerosis. Nat Med 24:1136–1142. 10.1038/s41591-018-0071-1

[58] Jackson KL, Dayton RD, Orchard EA, Ju S, Ringe D, Petsko GA, Maquat LE, Klein RL (2015) Preservation of forelimb function by UPF1 gene therapy in a rat model of TDP-43-induced motor paralysis. Gene Ther 22:20–28. 10.1038/gt.2014.101

[59] Cummings CJ, Sun Y, Opal P, Antalffy B, Mestril R, Orr HT, Dillmann WH, Zoghbi HY (2001) Over-expression of inducible HSP70 chaperone suppresses neuropathology and improves motor function in SCA1 mice. Hum Mol Genet 10:1511–1518.

[60] Chai Y, Koppenhafer SL, Bonini NM, Paulson HL (1999) Analysis of the role of heat shock protein (Hsp) molecular chaperones in polyglutamine disease. J Neurosci 19:10338–10347. 10.1523/jneurosci.19-23-10338.1999

[61] Wyttenbach A, Carmichael J, Swartz J, Furlong RA, Narain Y, Rankin J, Rubinsztein DC (2000) Effects of heat shock, heat shock protein 40 (HDJ-2), and proteasome inhibition on protein aggregation in cellular models of Huntington’s disease. Proceedings of the National Academy of Sciences of the United States of America 97:2898–2903. 10.1073/pnas.97.6.2898

[62] Laferrière F, Maniecka Z, Pérez-Berlanga M, Hruska-Plochan M, Gilhespy L, Hock EM, Wagner U, Afroz T, Boersema PJ, Barmettler G, Foti SC, Asi YT, Isaacs AM, Al-Amoudi A, Lewis A, Stahlberg H, Ravits J, De Giorgi F, Ichas F, Bezard E, Picotti P, Lashley T, Polymenidou M (2019) TDP-43 extracted from frontotemporal lobar degeneration subject brains displays distinct aggregate assemblies and neurotoxic effects reflecting disease progression rates. Nat Neurosci 22:65–77. 10.1038/s41593-018-0294-y

[63] Renkawek K, Bosman GICGM, de Jong WW (1994) Expression of small heat-shock protein Hsp27 in reactive gliosis in Alzheimer disease and other types of dementia. Acta Neuropathologica 87:511–519. 10.1007/BF00294178

[64] Oslowski CM, Urano F (2011) Measuring ER stress and the unfolded protein response using mammalian tissue culture system. Methods Enzymol 490:71–92. 10.1016/B978-0-12-385114-7.00004-0

[65] Hetz C (2012) The unfolded protein response: controlling cell fate decisions under ER stress and beyond. Nature Reviews Molecular Cell Biology 13:89–102. 10.1038/nrm3270

[66] Ron D, Walter P (2007) Signal integration in the endoplasmic reticulum unfolded protein response. Nature Reviews Molecular Cell Biology 8:519–529. 10.1038/nrm2199

[67] Hetz C, Mollereau B (2014) Disturbance of endoplasmic reticulum proteostasis in neurodegenerative diseases. Nature Reviews Neuroscience 15:233–249. 10.1038/nrn3689

